# PE/PPE proteins contribute to *Mycobacterium tuberculosis* drug resistance

**DOI:** 10.1101/2025.09.24.678375

**Authors:** Vishant Boradia, Junxi Chen, Andrew Frando, Lindsay V. Clark, Christoph Grundner

**Author notes:** Department of Medicine, University of Washington, Seattle, WA, USA.

## Abstract

The outer membrane (OM) of mycobacteria is a formidable permeability barrier that confers drug tolerance, and whether drugs traverse the OM by mechanisms beyond passive diffusion remains unclear. The proline-glutamic acid (PE) and proline-proline-glutamic acid (PPE) proteins of pathogenic mycobacteria include several OM transporters. Here, we tested the role of PE/PPE proteins in *Mycobacterium tuberculosis* (*Mtb*) drug transport and resistance. Mutations in multiple *pe/ppe* genes were strongly associated with drug resistance in a genetic association study, and mutations in *ppe42* and *ppe51* also conferred increased resistance *in vitro*. Deletion of a *pe/ppe* pair transcriptionally responsive to drug exposure, *pe25/ppe41*, led to elevated resistance to isoniazid (INH) across all major *Mtb* lineages and accelerated INH resistance emergence *in vitro*. These data identify a role for *Mtb* PE/PPE proteins in drug resistance consistent with the PE/PPE transporter paradigm, suggest a broader contribution of this large protein family, and a new factor of *Mtb* clinical drug resistance.

## INTRODUCTION

The outer membrane (OM) of mycobacteria consists of long-chain fatty acids that severely restrict solute diffusion ^1^. This OM barrier is also highly impermeable to many clinical drugs and at least partially responsible for the high intrinsic drug tolerance of *Mycobacterium tuberculosis* (*Mtb*) and other pathogenic mycobacteria ^2^. While non-pathogenic mycobacteria and gram-negative bacteria with a comparable OM express porins to facilitate free diffusion, *Mtb* and other pathogenic mycobacteria lack canonical porins. As a result, OM permeability of mycobacteria is typically two to three orders of magnitude lower than that of gram-negative bacteria ^3^ and poses one of the many challenges to effective tuberculosis (TB) treatment.

A unique feature of the genomes of pathogenic mycobacteria is the presence of two large gene families with mostly unknown function, the *pe* and *ppe* genes. *Mtb*, for example, encodes approximately 100 PE and 69 PPE proteins, which together take up nearly 10% of the genome’s coding capacity ^4^. Despite their abundance, a unifying or shared function for the PE/PPE proteins has not yet been discovered. Several recent studies now show that some PE/PPE proteins function as specific pores or channels in nutrient transport across the OM ^5-8^, and pore formation has now directly been shown for one PPE protein ^9^. This emerging transport function aligns with the relative occurrence of PE/PPE proteins and porins in mycobacteria: PE/PPE proteins are typically present in much larger numbers when porins are absent.

TB is exceedingly difficult to treat due to several host and bacterial factors. In addition to its impermeable OM, *Mtb* expresses inner membrane efflux pumps that effectively reduce intracellular drug concentrations ^10^. Efflux pumps are commonly upregulated in clinical drug-resistant strains and are a major contributor to *Mtb* drug tolerance and resistance to virtually all drugs currently in use ^11^. How drugs traverse the impermeable OM of pathogenic mycobacteria in either direction in the absence of porins, however, has remained an open question.

Here, we show that several PE/PPE proteins affect drug resistance in a way that is consistent with transporter function and are likely facilitating drug uptake across the OM. These data challenge the idea that new TB drugs need to be lipophilic to penetrate the outer membrane and show that the *pe/ppe* genes contribute to clinical drug resistance. With 169 *pe/ppe* genes in *Mtb*, these gene families could widely affect drug susceptibility and be a central contributor to TB drug resistance.

## RESULTS

### Mutations in *pe/ppe* genes are associated with clinical drug resistance

To identify candidate *pe/ppe* genes that may contribute to clinical drug resistance, we carried out a targeted genetic association study with a collection of 16,891 genome sequences of drug- and multi-drug resistant *Mtb*. Whole genome sequence data of *Mtb* was obtained from three sources: 2,659 genome sequences from the Wellcome Trust Sanger Institute pilot study ^12^, 3,922 from the NIH-NIAID TB Portals collection ^13^, and 10,310 from the CRyPTIC study ^14^. We analyzed sequences of the *pe* and *ppe* genes along with known resistance loci (*aphC, gyrA, inhA, katG*, and *rpoB*) as positive controls. Because of their high sequence variability and complex repetitive sequences, DNA sequence quality and genome assembly across the *pe*/*ppe* genes can be poor when using short-read sequencing, which is predominantly used for sequencing of drug resistant *Mtb*. As a result, many GWAS studies excluded *pe*/*ppe* genes from analysis. To assess data quality and possible bias, we analyzed the read depth and mapping quality across *pe, ppe*, and *pe-pgrs* genes from the set of nearly 4,000 genomes from the NIH collection. Read depth was generally comparable between pe/ppe and all other genes. Mapping quality for the *pe* genes was also comparable to all non-*pe*/*ppe* genes but was reduced for the *ppe* genes and more so for the *pe-pgrs* genes (Supplemental Fig. 1). The *pe-pgrs* subfamily is also known to harbor large sequence variation independent of drug resistance ^15-17^ and together with the *ppe-mptr* subfamily is the main source of errors in *Mtb* genome sequences ^18^. This variation and the reduced mapping quality led us to exclude the *pe-pgrs* genes from further analysis. We considered promoter and gene body mutations for the genetic association analysis. Using the linear mixed model approach in PySEER, some of the most significant associations were found with mutations in *ppe42, ppe35*, and *ppe51* (Fig. 1a). The *ppe42* mutation most significantly associated with drug resistance introduces a premature stop codon and was previously noted by the CRyPTIC consortium to be associated with amikacin and kanamycin resistance ^14^. Mutations in *ppe35* have also previously been linked to pyrazinamide resistance ^19,20^, and *ppe51* has experimentally been implicated in resistance to isoniazid (INH), although a transporter or channel role for PE/PPE proteins was not recognized at the time ^21^. Additional associations with drug resistance that had high odds ratios and low false discovery rate (FDR) were found in *ppe3, pe29*, and *ppe34-37* (Fig. 1a, Table 1). Further filtering to associations with FDR <0.00001, we identified 35 promoter and 443 coding variants associated with resistance to at least one drug within our target genes, in addition to the five positive controls (Supplemental Table 1).

**Table 1:** *pe* and *ppe* genes associated with drug resistance in the genetic association study. The ten mutations with the lowest FDR and odds ratio >10 are shown.

| **Gene** | **Drug** | **Variant** | **FDR global** | **p-value** | **Allele frequency** | **Odds ratio** |
| --- | --- | --- | --- | --- | --- | --- |
| *ppe42* | Amikacin | Y290* | 4.86E-48 | 1.94E-68 | 3.69E-03 | 116.36 |
| *ppe35* | Ethambutol | I539V | 1.35E-46 | 1.30E-56 | 7.31E-03 | 37.07 |
| *pe29* | Amikacin | G30R | 1.02E-41 | 2.88E-64 | 3.69E-03 | 84.67 |
| *ppe42* | Kanamycin | Y290* | 8.74E-40 | 5.45E-52 | 3.82E-03 | 126.06 |
| *pe29* | Kanamycin | G30R | 9.02E-35 | 5.84E-49 | 3.82E-03 | 81.76 |
| *ppe3* | Amikacin | G60E | 8.64E-33 | 6.79E-280 | 1.67E-02 | 86.47 |
| *ppe51* | Amikacin | A222T | 1.10E-31 | 5.87E-56 | 3.30E-03 | 74.39 |
| *ppe36* | Ethionamide | W125R | 8.77E-31 | 1.50E-62 | 1.03E-02 | 20.87 |
| *ppe37* | Amikacin | D381N | 1.65E-30 | 6.21E-278 | 1.68E-02 | 82.34 |
| *ppe47* | Amikacin | E124G | 2.63E-30 | 8.90E-278 | 1.66E-02 | 85.79 |

**Figure 1.**
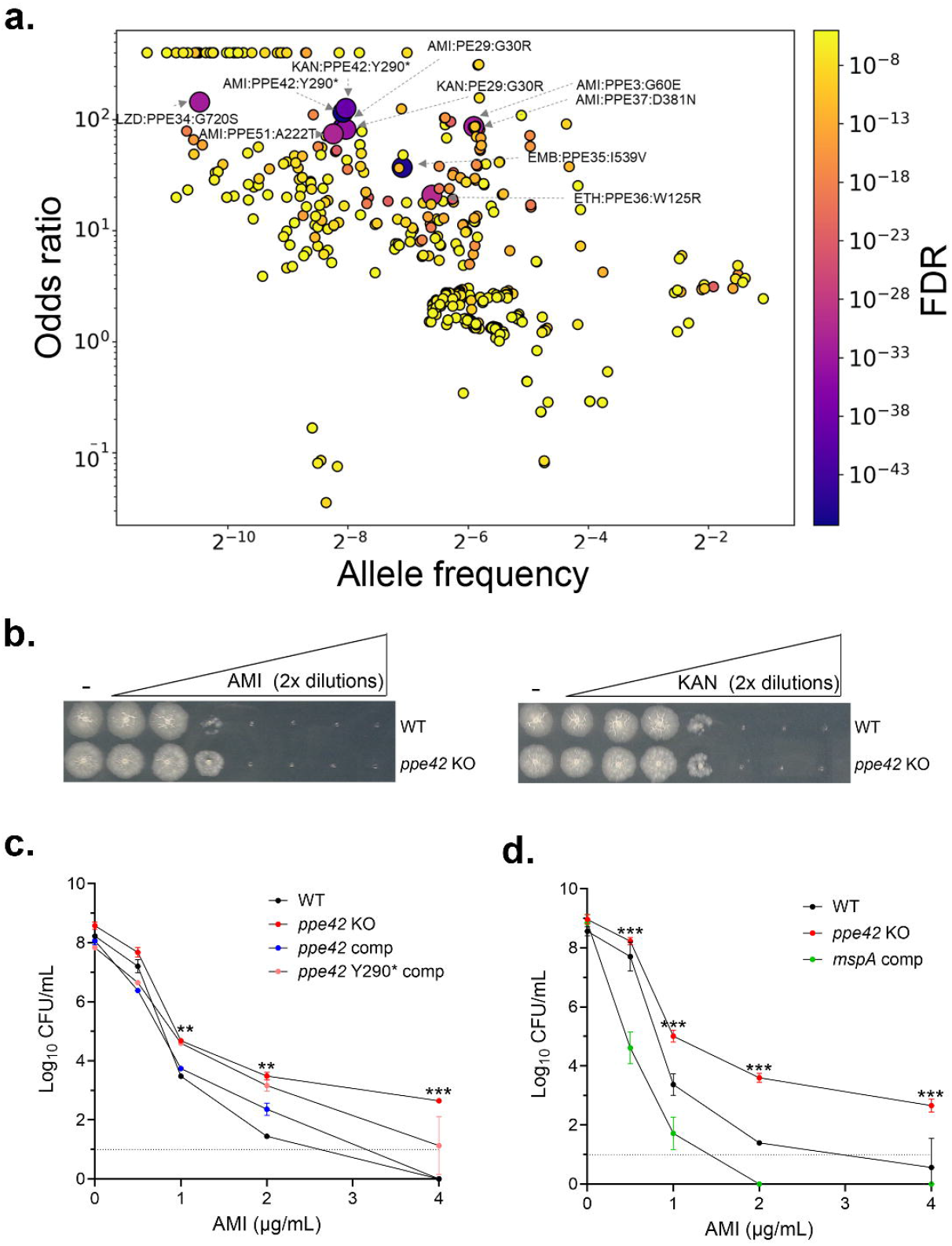
Multiple *pe/ppe* mutations are associated with clinical drug resistance. **a**. Genetic associations of *pe/ppe* mutations and clinical drug resistance are shown as scatterplot with odds ratio and allele frequency. False discovery rate (FDR) is shown according to color scheme (right). Larger symbols indicate the top ten candidates based on FDR, and the respective drug and mutation are indicated. *pe-pgrs* genes were excluded from the analysis. **b**. Spotting assay after exposure to drug shows that deletion of *ppe42* increases resistance to amikacin, but not kanamycin. Cultures were grown with drug for 7 days before spotting on agar without drug. **c**. Deletion of *ppe42* is partially complemented by WT *ppe42* but not the Tyr290 truncation, as shown by CFU assay. **d**. The MspA porin reverses the resistance of the *ppe42* KO to amikacin. Error bars for CFU assays show standard deviation of three biological replicates. P-values for comparisons in c. and d. are given in Supplemental Table 2. Abbreviations: AMI: amikacin. EMB: ethambutol. KAN: kanamycin. LZD: linezolid. We used Gemini and ChatGPT for writing a Python script to generate Fig. 1a. We reviewed and validated all AI outputs before inclusion.

### *ppe42* mutation leads to amikacin resistance

To test whether the *ppe42* mutation causes amikacin and/or kanamycin resistance, we generated a *ppe42* deletion strain by recombineering ^22^, and complemented this strain with wild-type (WT) and *ppe42* with the associated mutation, a stop codon at Tyr290. The complex outer membrane lipids Phthiocerol Dimycocerosates (PDIM) can be lost during *in vitro* culture and affect OM permeability ^8,23^. To maintain PDIM, we grew all cultures in 100µM sodium propionate ^23^. We incubated liquid culture of H37Rv WT and the *ppe42* deletion mutant in the presence of drugs for seven days and determined cell viability by spotting on solid medium. The *ppe42* deletion strain was more resistant to amikacin, a second-line drug used for treatment of multi-drug resistant tuberculosis ^20^, but not kanamycin when compared to WT (Fig. 1b). We next plated amikacin-treated strains on solid media for quantitating the difference in amikacin susceptibility by colony forming units (CFU) assay. At amikacin concentrations near the minimal inhibitory concentration (MIC), the *ppe42* KO mutant was more resistant, with ∼2log higher CFU than the WT. The *ppe42* deletion strain regained some sensitivity to amikacin when complemented with WT *ppe42*, but not when complemented with the copy of *ppe42* that carried the clinical mutation (Fig. 1c, see Supplemental Table 2 for all p-values, CFU data for kanamycin in Supplemental Fig. 2). Since the mutated *ppe42* phenocopies the deletion strain, the clinical mutation is likely a loss of function mutation. To test if the increased drug resistance of the *ppe42* deletion strain is related to transport across the OM, we next tested if the *M. smegmatis* porin MspA can functionally complement the *ppe42* deletion. MspA has previously been shown to be an OM porin in nonpathogenic mycobacteria that can bypass the outer membrane when introduced into pathogenic mycobacteria ^24^. To test if this OM porin can revert the phenotype of *ppe42* deletion, we expressed *mspA* in the *ppe42* knockout strain. Expression of *mspA* fully restored amikacin susceptibility in the *ppe42* deletion background (Fig. 1d), suggesting that *ppe42* is also an OM conduit. These data show that the truncation at Tyr290 in *ppe42* observed in clinical drug resistance confers drug resistance to amikacin but not kanamycin.

### *ppe51* mutation leads to resistance to multiple drugs

To experimentally test the association of *ppe51* with drug resistance, we generated a *ppe51* deletion strain and tested its susceptibility to drugs. Because the association of a genetic mutation especially with second-line drugs is often confounded by multiple drug-resistance, we tested against a panel of ten drugs currently in clinical use. The deletion mutant showed increased susceptibility to amikacin, capreomycin, moxifloxacin, streptomycin, and INH (Fig. 2a). Because deletion of *ppe51* conferred resistance to multiple drugs, we explored potential compensatory genetic effects of *ppe51* deletion by comparing global transcription in the WT and deletion strains by RNA-seq. Deletion of *ppe51* led to multiple significant changes in gene expression compared to WT (Fig. 2b). Notably, transcripts for six inner membrane efflux pump genes (*mmr, mmpl5, mmps5, Rv1216c-1218c*) were significantly induced (Fig. 2b, c, all differentially expressed genes in Supplemental Table 3). Since these genes are known to efflux TB drugs including INH ^11^, the effect of *ppe51* deletion on drug susceptibility is likely indirect and shows coupling of *ppe51* with inner membrane transport.

**Figure 2.**
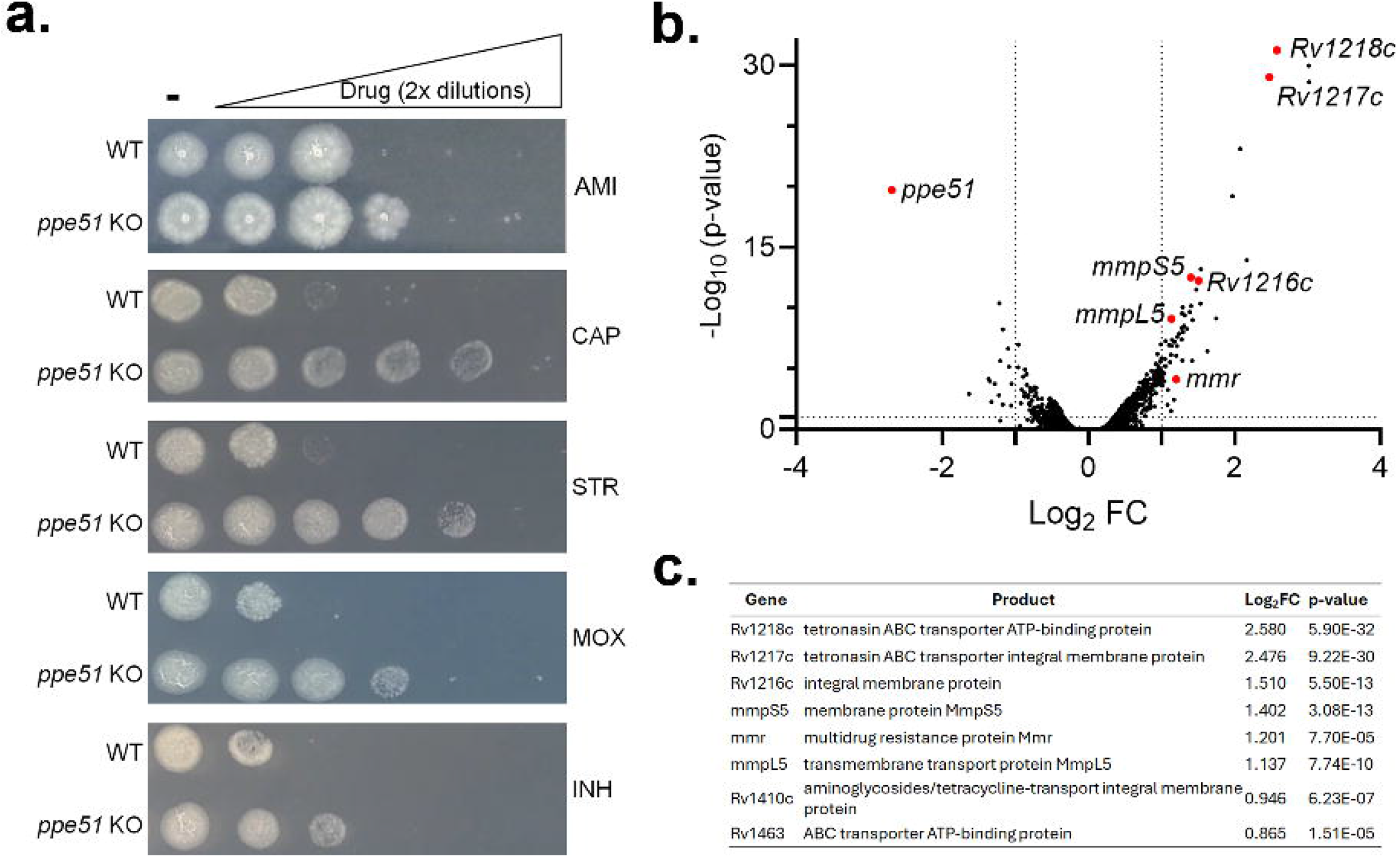
Deletion of *ppe51* increases drug resistance and upregulates inner membrane transporters. **a**. Spotting of WT and *ppe51* KO strain grown in presence of drugs in 2-fold dilution shows increased resistance to multiple drugs. **b**. RNA-seq analysis shows increased transcription of several known efflux pump genes as a result of *ppe51* deletion. **c**. The efflux pump genes affected by *ppe51* deletion. All differentially expressed genes are given in Supplemental Table 3. Abbreviations: AMI: amikacin. CAP: capreomycin. STR: streptomycin. MOX: moxifloxacin. INH: isoniazid.

### *ppe3, ppe35*, and *ppe36* deletion do not affect drug resistance

We tested additional candidates from our association study for effects on drug resistance. We tested a *ppe35* deletion strain for resistance to pyrazinamide or any other of ten clinical drugs, but the deletion strain did not alter the sensitivity to drugs. Similarly, deletion of *ppe3* and *ppe36* did not affect drug resistance to any of the ten tested drugs (Supplemental Fig. 2, 3). These data show the limitations of genetic association studies, which might be particularly challenging for the *pe-pgrs* and *ppe* genes due to the poorer mapping quality of sequencing data. These data also show that deletion of *ppe* genes does not generally affect the OM permeability but only does so in the case of specific *ppe* genes and in the context of specific drugs.

### *pe25/ppe41* affect susceptibility to INH

We next sought to identify additional candidates by an orthogonal approach. Transporters can respond transcriptionally to their substrates, a behavior that has also been reported for several *pe/ppe* transporters ^6,8^. To identify additional *pe/ppe* genes that might affect drug resistance but were not detected by our genetic association study, we mined publicly available RNA-seq data for changes in *pe/ppe* gene expression that is induced by drugs. Twenty-eight datasets from public repositories captured the transcriptional responses to ten drugs. From these data, we identified the *pe/ppe* genes that were significantly regulated in response to drug exposure and plotted them against the number of drugs to which they responded (Fig. 3a). Many *pe/ppe* transcripts responded to drug exposure, and several to different drugs. Generally, more *pe/ppe* transcripts were down-than upregulated in response to drugs (Fig. 3a). *ppe51* was also differentially regulated by several drugs, and *pe25/ppe41* showed the most consistent directionality of expression change across different drugs. To test if these candidates have any functional consequence for drug resistance, we generated deletion strains for five and tested their susceptibility with the ten-drug panel by spotting assay. The *pe25/ppe41* deletion strain was more resistant to INH than WT. None of the other deletion strains showed altered susceptibility to the drugs tested (Supplemental Fig. 4). To quantitate the effect of *pe25/ppe41* deletion on drug resistance, we determined the MIC of the WT and deletion strains by CFU assay. At concentrations of INH corresponding to the MIC of drug-susceptible *Mtb* (0.225-0.45µM), the *pe25/ppe41* deletion strain was more resistant than the WT strain, with >10-fold higher CFU and ∼2-fold increase in the MIC_50_ (Fig. 3b). Complementation of the *pe25/ppe41* deletion strain fully restored the susceptibility to INH (Fig. 3b).

**Figure 3.**
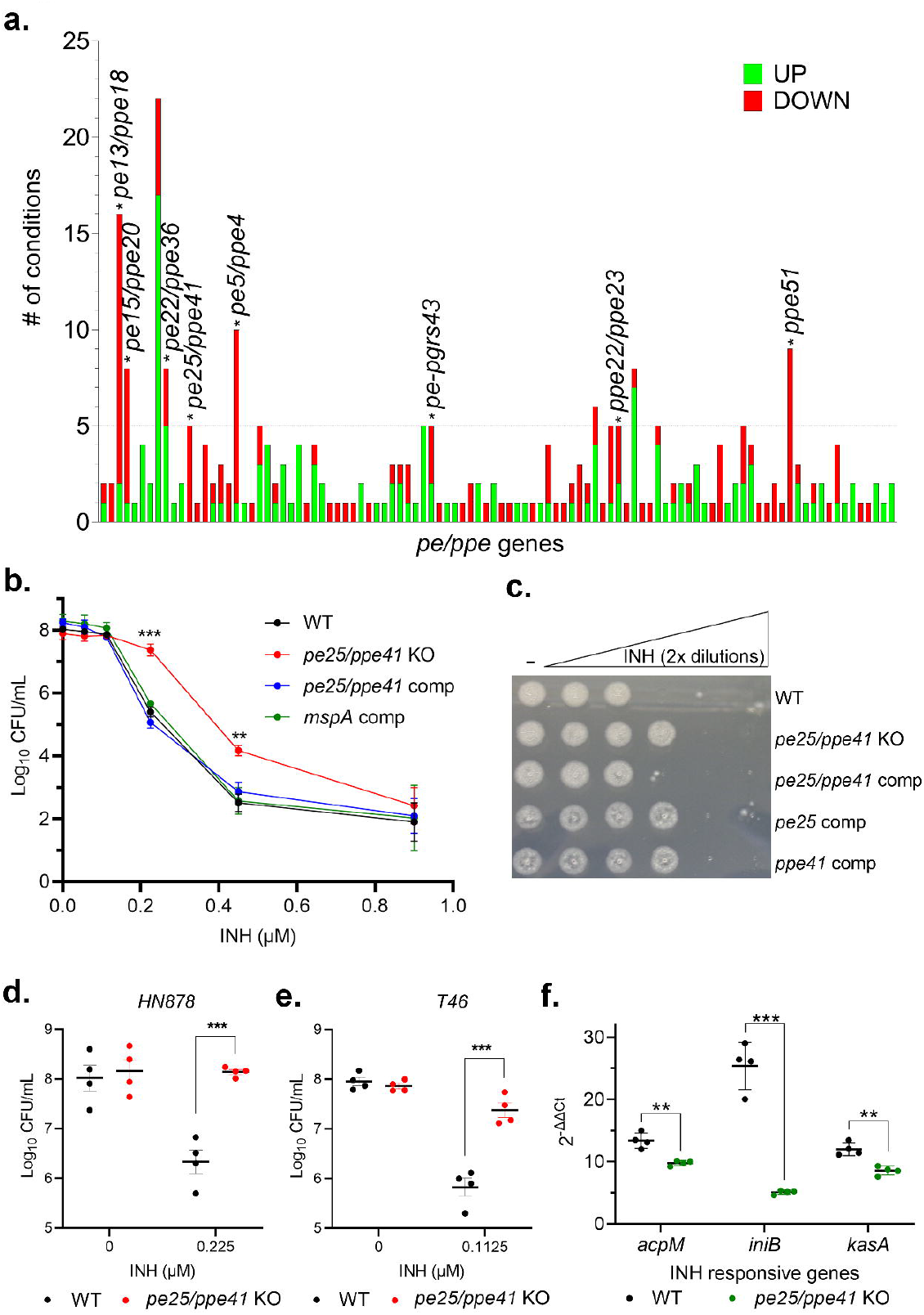
The PE25/PPE41 complex affects isoniazid susceptibility. **a**. The transcriptional response of *pe/ppe* genes to drugs. *pe/ppe* genes showing transcriptional up-(green) or downregulation (red) in response to drugs in historical RNA-seq data were plotted against the number of drugs they were found to respond to. Deletions of *pe* and *ppe* genes tested in this study are labeled. Direction of change is indicated by color. **b**. A *pe25/ppe41* deletion strain shows higher resistance to INH. The heterologous OM porin MspA from *M. smegmatis* can functionally complement the *pe25/ppe41* deletion. Bacteria were enumerated by colony forming units (CFU) assay and data are shown as the mean ± SEM of two experiments performed in quadruplicate. Statistical significance was determined by multiple comparison t-test. **: p< 0.01, ***: p< 0.001. **c**. Complementation of the *pe25/ppe41* deletion restores susceptibility to INH, but not complementation with either *pe25* or *ppe41* alone, as shown by spotting assay. **d**. Deletion of *pe25/ppe41* also reduces the susceptibility to clinically relevant concentrations of INH in *Mtb* strains from lineage 1 (T46) and **e**. lineage 2 (HN878) **f**. The known INH-responsive genes *iniB, acpM*, and *kasA* are induced by INH exposure in the H37Rv WT but less so in the *pe25/ppe41* deletion strain, as determined by qRT-PCR. Datapoints show the average of two technical replicates for four biological replicates **: p< 0.01, ***: p< 0.001.

To test if the increased drug resistance of the *pe25/ppe41* deletion strain is related to transport across the OM, we next tested if the *M. smegmatis* porin MspA can functionally complement the *pe25/ppe41* deletion. MspA restored the susceptibility of the *pe25/ppe41* deletion strain to INH to WT levels, suggesting that PE25/PPE41 has a similar function to MspA in transport across the OM (Fig. 3b). The decreased susceptibility to INH upon *pe25/ppe41* deletion indicates that the PE25/PPE41 complex imports INH. To test if *pe25, ppe41*, or both affect INH sensitivity, we complemented the *pe25/ppe41* deletion strain with each gene individually under the control of an anhydrotetracycline (ATc)-inducible promoter. Complementation with both but not with *pe25* or *ppe41* alone restored sensitivity to INH, showing that both are required (Fig. 3c). The genetic background of a strain can affect drug resistance phenotypes ^25,26^. To test the effect of strain background and whether the *pe25/ppe41* effect on INH susceptibility is conserved beyond the lineage 4 strain H37Rv, we introduced *pe25/ppe41* deletions in representative strains from the major circulating lineages—T46 (lineage 1) and HN878 (lineage 2). Both strains showed similar changes in INH resistance to the lineage 4 reference strain H37Rv (Fig3d, e).

### Loss of *pe25/ppe41* reduces transcription of INH-responsive genes

INH is a prodrug that requires activation by KatG in the cytoplasm, and it has well-defined transcriptional effects ^27^. To further test for an INH import function of *pe25/ppe41* and to test whether *pe25/ppe41* alone can alter the levels of INH in the cytoplasm, we tested if *pe25/ppe41* deletion affects the expression of INH-responsive genes. We chose the three INH-responsive genes *iniB, acpM*, and *kasA*, and compared their expression by qRT-PCR in WT and the *pe25/ppe41* deletion strain after treatment with 0.2µg/mL INH for 5 hours. INH strongly induced the expression of the three INH-responsive genes in WT but significantly less in the *pe25/ppe41* deletion strain (Fig. 3f). These data indicate that *pe25/ppe41* deletion is sufficient to reduce INH concentrations in the cytoplasm, which is consistent with an importer function.

### *pe25/ppe41* mutations accelerate drug resistance

Although neither *pe25* nor *ppe41* mutations were significantly associated with drug resistance in our genetic association analysis, anecdotal data also link *pe25/ppe41* to clinical drug resistance. The Lisboa cluster of *Mtb* strains has been associated with INH resistance and high rates of multidrug-resistant and extensively drug-resistant tuberculosis in Portugal ^28-30^. Although INH resistance in these strains is associated with two canonical *inhA* mutations ^31^, strains in the Lisboa3 cluster also carry a truncation of *ppe41* after bp112 ^29^. To test if this *ppe41* truncation could also contribute to INH resistance, we complemented the H37Rv *pe25/ppe41* deletion strain with a WT or truncated copy of *ppe41*. In contrast to the WT gene, the truncated *ppe41* did not restore INH sensitivity of the *pe25/ppe41* deletion strain (Fig. 4a), suggesting that it contributes to high-level INH resistance in the Lisboa3 strain and/or promotes the emergence of canonical INH mutations. To quantitate the effect of the truncation, we determined the CFU of the WT strain and a *pe25/ppe41* deletion strain complemented with the truncated *ppe41*. The truncated ppe41 conferred higher resistance, with ∼2 orders of magnitude higher CFUs (Fig. 4b). To test the idea that *pe25/ppe41* deletion can affect the rate of emergence of INH resistance, we plated cultures of WT and the *pe25/ppe41* deletion strain on agar plates containing INH at 10-, 25-, 125-, and 500-fold the MIC and quantified the resistant colonies that grew under drug pressure by CFU. The *pe25/ppe41* deletion strain developed INH-resistant colonies at an about 3.5-fold higher rate than the WT strain across all tested concentrations (Fig. 4c). Sequencing of resistance mutants showed that resistance in the *pe25/ppe41* deletion strain was due to canonical *katG* mutations typically associated with INH resistance *in vitro* (Supplemental Fig. 5). These findings indicate that loss of *pe/ppe* function accelerates the emergence of INH resistance.

**Figure 4.**
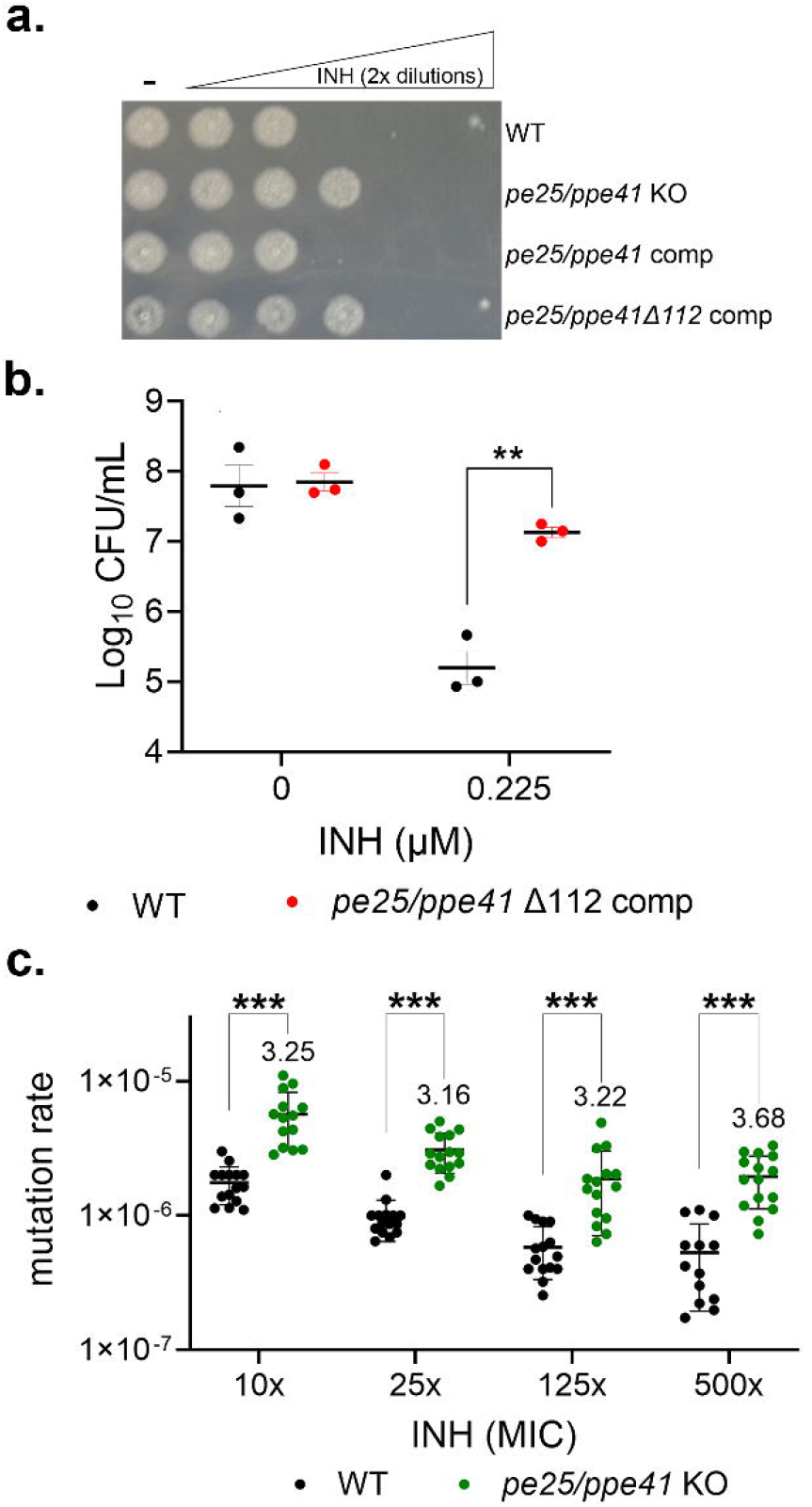
*ppe41* truncation contributes to INH resistance. **a**. The *ppe41* deletion at bp112 in the Lisboa3 strain family increases resistance of *Mtb* to INH, as shown by spotting after growth on drug for 7 days. **b**. *ppe41* truncation increases INH resistance as quantified by CFU assay. Data points represent average of two technical replicates for three biological replicates **c**. Deletion of *pe25/ppe41* increases the rate at which INH resistance emerges. *Mtb* cultures were plated onto 7H10+GO plates without drug and with INH at concentrations 10x–500x above the MIC. Colonies on drug-free plates represented the total viable cell count, while colonies on plates with drug were counted as resistant mutants. The mutation rate was calculated by dividing the number of colonies on drug-containing plates by the total viable cell count from non-drug plates plates. Each datapoint represents a biological replicate. Error bars are mean ± SD. Statistical significance was determined by multiple comparison t-test. ***: p< 0.001.

## DISCUSSION

*Mtb* and related pathogenic mycobacteria have some of the thickest and most impermeable outer membranes known in microbiology ^1,32^. The mechanism by which *Mtb* takes up nutrients through this thick barrier in the absence of canonical porins was a longstanding question until the first *Mtb* transporters from the PE/PPE protein families were discovered. The PE/PPE proteins are an idiosyncratic protein family with no apparent orthologs outside of mycobacteria, and they are generally more frequent in pathogenic mycobacteria that do not have canonical porins. Indeed, several PE/PPE proteins have now been shown to be nutrient transporters ^6,8^. These and data showing direct pore-forming function ^9^ further support the idea that OM transport might be a common function among the PE/PPE proteins.

Drug uptake by *Mtb* is generally presumed to occur by passive diffusion, an idea that has led to a guiding principle in TB drug development that TB drugs should be lipophilic. However, many effective TB drugs in clinical use are not. The effects of PPE42 on amikacin susceptibility, of PPE51 on several drugs, and of PE25/PPE41 on INH susceptibility are indeed consistent with an importer function, and the presence of proteinaceous OM drug importers would provide a plausible explanation for this contradiction. The effects of the PE/PPE proteins tested here on drug susceptibility are highly specific. Of nine PE/PPE proteins tested, only three showed altered drug susceptibility, two of which only to one drug. These data suggest that PE/PPE deletion does not cause a generalized permeability change in the OM but has specific effects on specific compounds.

In addition to mutations in *ppe42* and *ppe51*, loss of function mutations of *pe25/ppe41* could be a facile mechanism for INH tolerance and/or resistance, and our data indeed show a higher rate of emergence of INH resistance in the *pe25/ppe41* deletion strain *in vitro*, suggesting it could be a steppingstone mutation to high-level drug resistance, for example in some Lisboa strains. The contribution of *pe/ppe* genes to clinical drug resistance combined with the large number of PE/PPE proteins in *Mtb* represent a previously unrecognized factor of TB drug resistance and raise the possibility that PE/PPE proteins contribute to drug uptake (and perhaps efflux) more widely. Several observations indeed point to a larger role of PE/PPE proteins in drug susceptibility and resistance. A genome-wide chemical-genetic screen recently identified several *pe/ppe* genes that conferred increased or decreased susceptibility to drugs *in vitro* ^33^. These data are consistent with our findings for *pe25/ppe41* and INH but require experimental validation for others since CRISPRi is subject to polar effects, and expression of *pe/ppe* genes can be linked to expression of inner membrane transporters—as we have shown for *ppe51*. Also, the large number of *pe/ppe* genes makes redundancy likely, potentially masking the effects of single gene deletion or knockdown and complicating the identification of PE/PPEs that affect drug susceptibility and resistance.

The physiologic substrate(s) and cellular function(s) of PPE42 and PE25/PPE41 remain to be identified. Although drugs are arguably accidental substrates of PE/PPE transporters, their cellular substrates might bear chemical resemblance to the drugs they transport. Interestingly, PPE42 contains a predicted serine hydrolase domain. Serine hydrolases play an important role in the resistance to beta-lactams (such as amikacin and kanamycin) through drug inactivation. The predicted AlphaFold structure for the PPE42 hydrolase domain shows a canonical catalytic site organization and a Ser residue in the expected position, suggesting it might be an active enzyme. However, loss of the hydrolase domain in the *ppe42* clinical resistance strain would have the opposite effect to known hydrolase-mediated drug resistance mechanisms and not lead to drug inactivation. The molecular function of the hydrolase domain in amikacin resistance needs further study, but it could serve as a binding domain for drug and thus facilitate entry. Our data identify a function for PE/PPE proteins in drug susceptibility that is consistent with their emerging OM transporter function. Although transporter function of PE/PPE proteins has now been shown in several cases, and many previously observed PE/PPE phenotypes are fully consistent with transporter function ^7,34^, much remains unknown about these most idiosyncratic protein families. For example, the topology and composition of the complexes, which are and which are not pore-like, the way in which pores or channels are formed across the OM, and the biochemical mechanism of transport all remain unknown but will inform drug development, drug targeting, diagnosis of drug resistance, and new strategies to counter drug resistance.

## ACKNOWLEGEMENTS

This work was supported by National Institutes of Health (NIH) grant R01AI180452 to CG.

## MATERIALS AND METHODS

### Genetic association analysis

Sequences were aligned to the MT_H37R7_V3 reference sequence using BWA-MEM ^35^, and the assembled genomes were aligned to the same reference using Minimap2 ^36,37^. The 17,047 files were then processed by BCFtools mpileup and BCFtools call ^38^ to make a joint-called VCF with haploid genotype calls. We considered promoter and gene body mutations for the analysis. A selection of 7,402 SNPs with less than 10% missing data in all three datasets and a minor allele count of at least 5, representing a random 10% of genomic regions, were used for estimating relatedness among strains. Relatedness was estimated by multiplying the centered genotype matrix by its transpose. Individual relatedness values were divided by the number of non-missing variants in common between two samples, and negative relatedness values were set to zero. The lineage of each strain was identified based on the set of lineage-specific SNPs packaged with TBprofiler v6.5.0 ^39^. Samples were retained if they had less than 10% missing data, showed agreement between assigned lineage and relatedness to other strains, and had antibiotic resistance phenotype data available, leaving 16,891 strains. We tested the 104 *pe/ppe* genes for association with phenotypic drug resistance. We excluded the *pe-pgrs* subfamily (64 genes) because it is known to have highly variable sequence. We included *aphC, gyrA, inhA, katG*, and *rpoB* as controls with known resistance associations. Within these 109 genes, we identified 19,154 nucleotide variants after splitting multiallelic variants into individual biallelic variants. Association analysis was performed at the nucleotide level for promoter variants and amino acid level for coding variants. Coding variants were agglomerated by amino acid position and type of variant (substitution, frameshift, in-frame insertion or deletion). Any given promoter or coding variant was used in association analysis with a given drug resistance trait if it had at least five strains with the minor allele and a recorded phenotype, totaling 357 unique promoter variants and 3,418 unique agglomerated coding variants across all 25 drugs. Association testing was conducted with PySeer ^40^ using the linear mixed model approach, with the relatedness matrix as a random effect (similarity matrix), and dataset (Sanger Institute, CRyPTIC, TB Portals) as a fixed covariate. At FDR<0.00001 across all tests, 35 promoter variants and 478 coding variants were significant for resistance to at least one drug. If the number of susceptible individuals with the minor allele was required to be zero, and the minor allele had to appear in resistant strains of at least two major *Mtb* lineages, there were six promoter variants and 41 coding variants significant for resistance to at least one drug within our target genes, including the five positive controls and 11 out of the 104 *pe/ppe* genes.

### Media and growth conditions

*Mycobacterium tuberculosis (Mtb*) H37Rv, a lineage 4 strain, was used as the parental strain for generating all mutants unless indicated otherwise. Strains were grown in Middlebrook 7H9 medium (Difco), supplemented with 10% (vol/vol) oleic acid-albumin-dextrose-catalase (OADC) enrichment (BBL; Becton Dickinson), 0.5% glycerol, and 0.05% Tween 80 or 0.05% Tyloxapol. The medium is referred to as “7H9+GO” when supplemented with glycerol and OADC only, “7H9+GOT” with Tween 80, “7H9+GOTy” with Tyloxapol, and “7H9+GOTyP” with the addition of 100 µM sodium propionate to maintain PDIM levels ^23^. For solid media, 7H10 agar was supplemented with 10% OADC and 0.5% glycerol. Strains harboring antibiotic resistance cassettes were cultured with the appropriate antibiotic: 50 μg/mL hygromycin, 30 μg/mL kanamycin, or 25 μg/mL zeocin. All drugs used for the drug screening assay were purchased from Sigma, and stock solutions were prepared in either water or DMSO.

### Creation of deletion and complemented strains

The deletion strains were generated using recombineering, as described previously ^22^. Initially, 500 bp upstream and downstream of the gene of interest, along with the hygromycin resistance cassette (primers Hyg_F and Hyg_R), were amplified separately (all primers used for each deletion are listed in Supplemental Table 4). The PCR fragments were then Gibson ligated to the 5′ and 3′ ends of the hygromycin resistance cassette to create the recombineering cassette. This linear recombineering cassette was PCR-amplified, purified, and electroporated into the *Mtb* H37Rv strain carrying the recombineering plasmid pNIT:Etc ^41^. Hygromycin-resistant colonies were screened, and the positive clones were confirmed by DNA sequencing. The PE25/PPE41 deletion was also generated in *Mtb HN878* and *T46* using the same recombineering pNIT:Etc strain and cassette used for *H37Rv*.

For the complemented strains, the *ppe42, ppe42 Y290*, pe*25, *ppe41, pe25/ppe41*, and *pe25/ppe41* (Δ112) coding regions, as well as the *M. smegmatis mspA* coding region were amplified by PCR using primers listed in Supplemental Table 4 and Gibson cloned into the pDTCF plasmid (Zeocin resistance) with a C-terminal FLAG tag, under the control of an anhydrotetracycline (ATc)-inducible promoter. The resulting plasmids were electroporated into the H37Rv *pe25/ppe41* deletion strain, and positive clones were selected by growth in hygromycin and zeocin.

### Drug screening

Drug screening was carried out in 96-well plates. Cultures were grown to log phase in 7H9+GOTyP media and incubated with drugs in 2-fold dilutions at a final optical density (OD) of 0.005. For pyrazinamide, we used medium at pH6. The assay plates were incubated at 37°C for 7 days, and cell viability was measured using either the agar spotting method, Alamar Blue assay, or colony-forming unit (CFU) counting. For the spotting method, cultures were mixed and 3 µL were spotted onto 7H10+GO agar plates, followed by incubation at 37°C for 2-3 weeks. For CFU counting, cultures were diluted and plated onto 7H10 agar plates, which were incubated for 3-4 weeks at 37°C.

### RNA sequencing

H37Rv and the *ppe51* deletion strain were grown to an OD_600_ of 0.8 in 7H9+GOTyP medium. Cells were pelleted and resuspended in buffered water, then incubated for an additional 5 hours. Following incubation, cells were pelleted at 4,000 g for 5 minutes at 4°C, resuspended in Trizol, and lysed by bead beating for 30 seconds at 6 m/s for 3 cycles, with intermittent cooling on ice. Cell debris was pelleted at 20,000 g for 1 minute, and the supernatant was transferred to a heavy phase-lock gel tube containing 300 µL of chloroform. The tubes were inverted for 2 minutes and centrifuged at 20,000 g for 5 minutes. RNA in the aqueous phase was precipitated with 300 µL of isopropanol and 300 µL of high-salt solution (0.8 M sodium citrate, 1.2 M sodium chloride). RNA was purified using the QIAGEN RNeasy kit, and ribosomal RNA was depleted using a previously published protocol ^42^. Briefly, a biotinylated oligo mixture of 23S, 16S, and 5S was incubated with RNA to anneal to rRNA, followed by incubation with streptavidin beads. mRNA was purified from the supernatant using Ampure XP beads. A cDNA library was generated using the NEBNext Ultra II RNA Library Prep Kit, and each replicate was barcoded in the DNA library using the NEBNext Multiplex Oligos for Illumina. Libraries were quantified using the KAPA qPCR quantification kit, pooled, and sequenced at the University of Washington Northwest Genomics Center using the Illumina NextSeq 500 High Output v2 Kit.Read alignment was performed using the Bowtie 2 custom processing pipeline (https://github.com/robertdouglasmorrison/DuffyNGS, https://github.com/robertdouglasmorrison/DuffyTools). Gene expression changes were identified using a combination of five differential expression tools within DuffyTools: round robin, RankProduct, significance analysis of microarrays (SAMs), EdgeR, and DeSeq2. The results from each DE tool were combined using a weighted average of fold change and significance (*p-value*). Genes with an averaged absolute fold change of more than 2-fold and a *p-value* <0.01 were considered differentially expressed.

### qRT-PCR analysis

Liquid cultures of *Mtb* H37Rv WT and *pe25/ppe41* deletion strains were grown to early log phase in 7H9+GOTyP media and subsequently treated with 0.2 µg/mL isoniazid for 5 hours. RNA was extracted using Trizol, purified, and cDNA was synthesized with SuperScript IV polymerase and random hexamer primers. mRNA expression levels for *iniB, acpM*, and *kasA* genes were quantified by qRT-PCR using SybrGreen iTaq chemistry. Expression levels were normalized to *sigA* mRNA expression, and the relative mRNA levels were calculated using the 2^− ΔΔCq^ method and plotted as the ratio of mRNA expression for each strain.

### Measuring the *in vitro* frequency of drug resistance

To measure the frequency of drug resistance, *Mtb H37Rv* WT and *pe25/ppe41* deletion strains were grown to an OD600 ∼0.6 in 7H9+GOTyP medium. The cultures were diluted to approximately 1 x 10^8^ and 1 x 10^7^ CFU/mL and plated onto non-drug-containing 7H10+GO agar plates to determine the total viable cell count, and drug-containing 7H10 agar plates supplemented with isoniazid at concentrations 10X–500X above the MIC. Plates were incubated at 37°C for 3-4 weeks. Colonies on drug-free plates represented the total viable cell count, while colonies on drug plates indicated resistant mutants. The frequency of resistance was calculated by dividing the number of colonies on drug-containing plates by the total viable cell count from non-drug plates.

## EXTENDED AND SUPPLEMENTARY DATA

**Supplemental Figure 1: Mapping quality comparison of *pe, ppe*, and *pe-pgrs* genes**. Within 4,031 protein-coding genes in the *M. tuberculosis* H37Rv v3 reference genome, 35 were classified as *pe*, 69 as *ppe*, 74 as *pe-pgrs*, and 3,863 as others (x-axis). Sequences from the 2,659 strains from the Wellcome Trust Sanger Institute pilot study were aligned to the reference using Bowtie2 and genotyped using GATK HaplotypeCaller, with joint calling performed by GATK GenomicsDBImport and GenotypeGVCFs. With each gene, the mapping quality (MQ) as reported by GATK was averaged across all variants (y-axis).

**Supplemental Figure 2: CFU data for WT, *ppe42* deletion, and *ppe35* deletion mutants grown in kanamycin and pyrazinamide, respectively**.

**Supplemental Figure 3: Negative spotting data for hits from genetic association study**. Cultures were grown with drug for 7 days before spotting on agar without drug.

**Supplemental Figure 4: Negative spotting data for candidates from the RNA-seq analysis**. Cultures were grown with drug for 7 days before spotting on agar without drug.

**Supplemental Figure 5: *katG* mutations in INH-resistant clones of the *pe25/ppe41* deletion strain**. Mutations were identified by DNA sequencing of the *katG* gene. The corresponding amino acid changes in KatG are shown. Numbers in parentheses indicate multiple occurrences of the mutation.

**Supplemental Table 1: Genetic associations between clinical drug resistance and *pe* and *ppe* genes**. All associations from our targeted association study with an FDR below 10^-5^ are listed.

**Supplemental Table 2: p-values for the CFU comparison of *ppe42* WT and mutant strains**. NS: Not significant. **: p<0.01. ***: p<0.001

**Supplemental Table 3: Differentially expressed genes in the *ppe51* deletion strain compared to WT**. RNA-seq analysis was done after four days of growth.

**Supplemental Table 4: Primers used for generating knockout strains, complementation strains, and primers used for qRT-PCR**.

