## Supplemental Table 2 for "PE/PPE proteins contribute to *Mycobacterium tuberculosis* drug resistance"

| **Fig 1c** | | |
| --- | --- | --- |
| **AMI (µg/mL)** | **Comparison** | **significance** |
| 1 | *ppe42* KO vs WT | ** |
|  | *ppe42* KO vs *ppe42* comp | *** |
|  | *ppe42* KO vs *ppe42 Y290** comp | NS |
| 2 | *ppe42* KO vs WT | ** |
|  | *ppe42* KO vs *ppe42* comp | ** |
|  | *ppe42* KO vs *ppe42 Y290** comp | NS |
| 4 | *ppe42* KO vs WT | *** |
|  | *ppe42* KO vs *ppe42* comp | *** |
|  | *ppe42* KO vs *ppe42 Y290** comp | NS |
| **Fig 1d** | | |
| 1 | *ppe42* KO vs WT | ** |
|  | *ppe42* KO vs *mspA* comp | *** |
| 2 | *ppe42* KO vs WT | *** |
|  | *ppe42* KO vs *mspA* comp | *** |
| 4 | *ppe42* KO vs WT | * |
|  | *ppe42* KO vs *mspA* comp | *** |
