## Supplementary figures and images for "PE/PPE proteins contribute to *Mycobacterium tuberculosis* drug resistance"

### Supplemental Figure 1

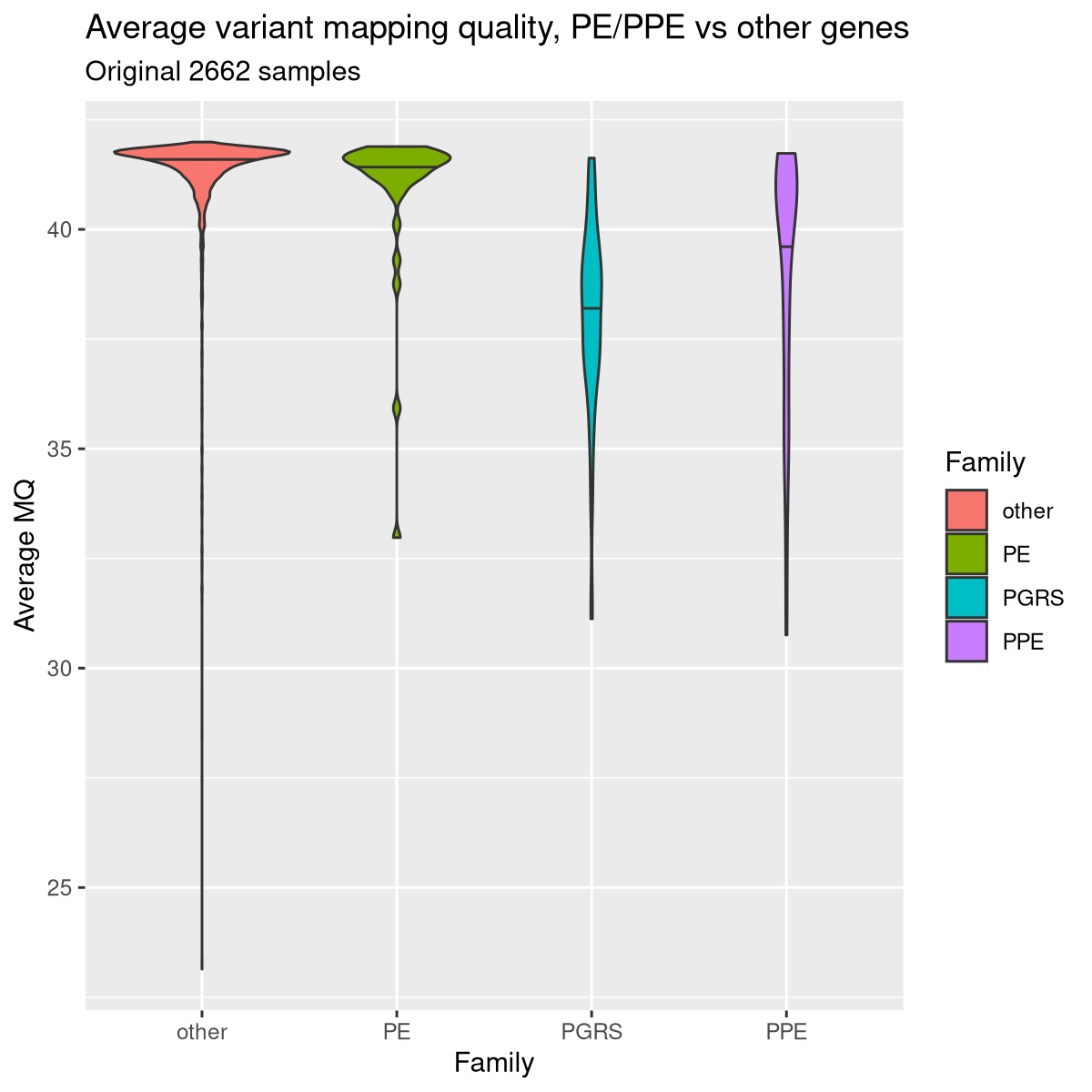

### Supplemental Figure 2

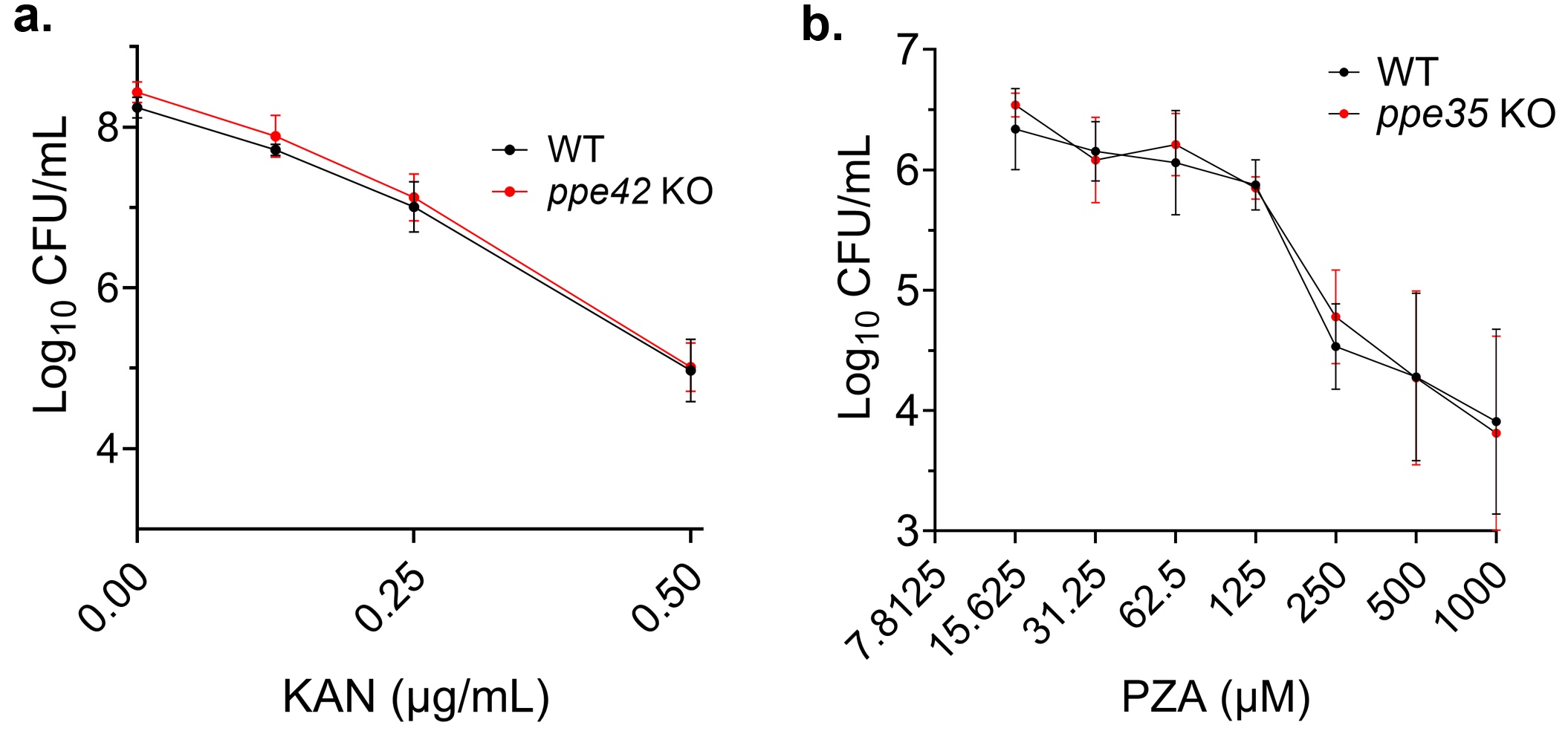

### Supplemental Figure 3

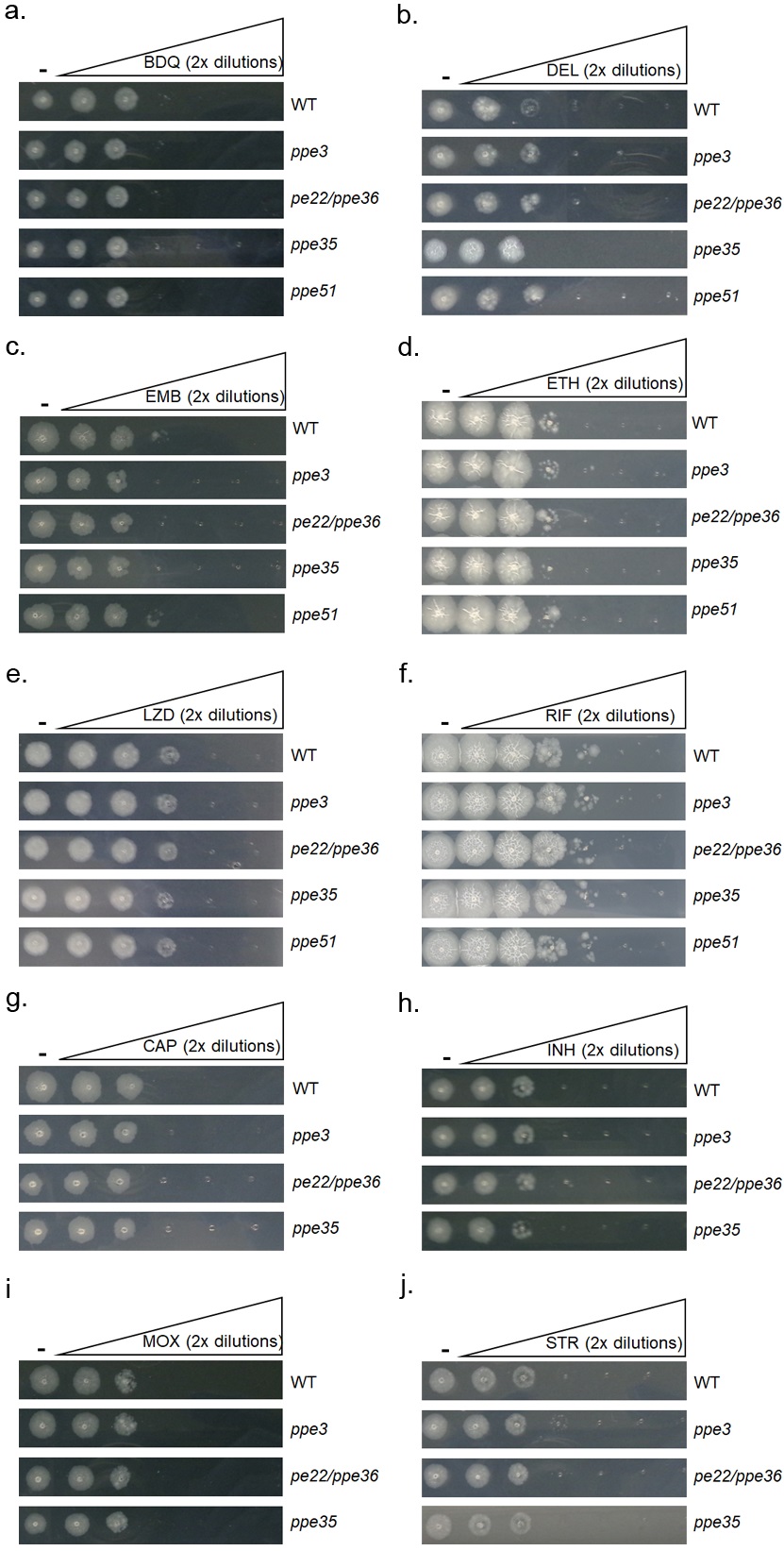

### Supplemental Figure 4

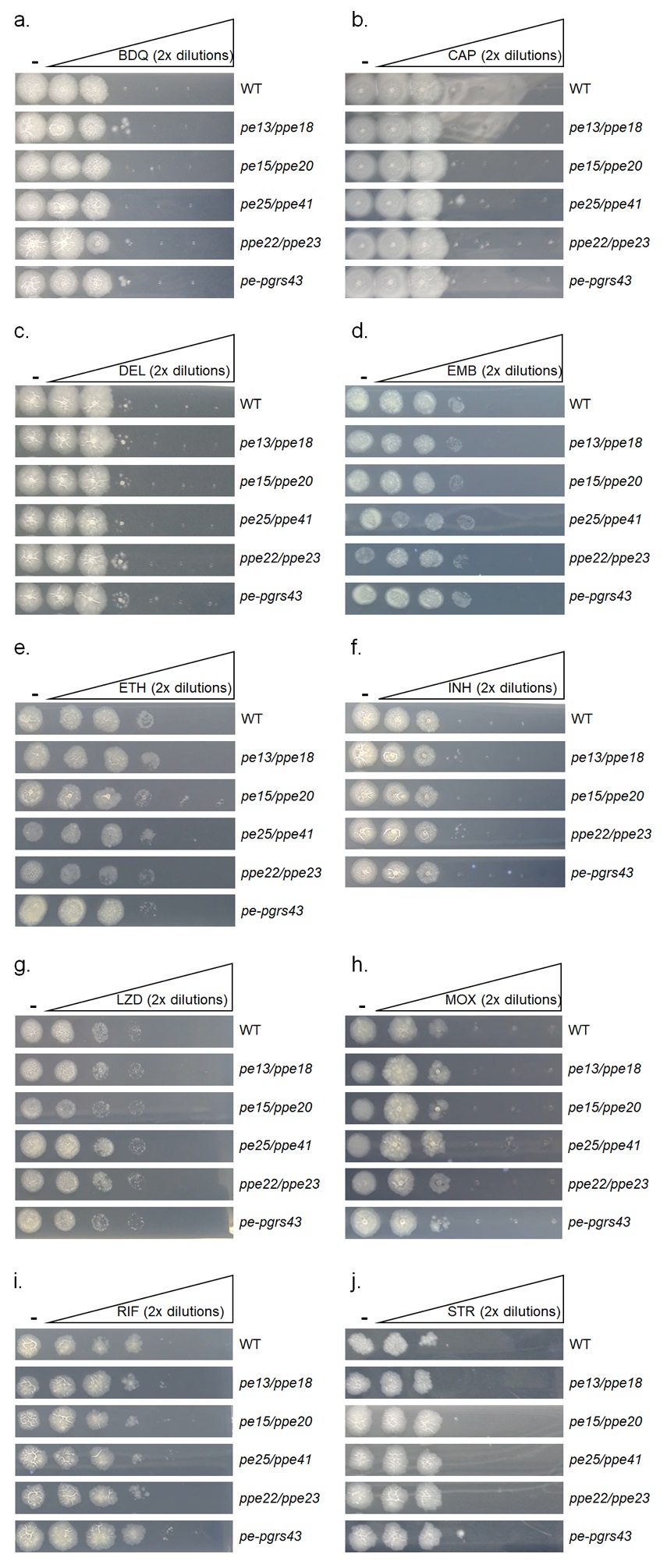

### Supplemental Figure 5

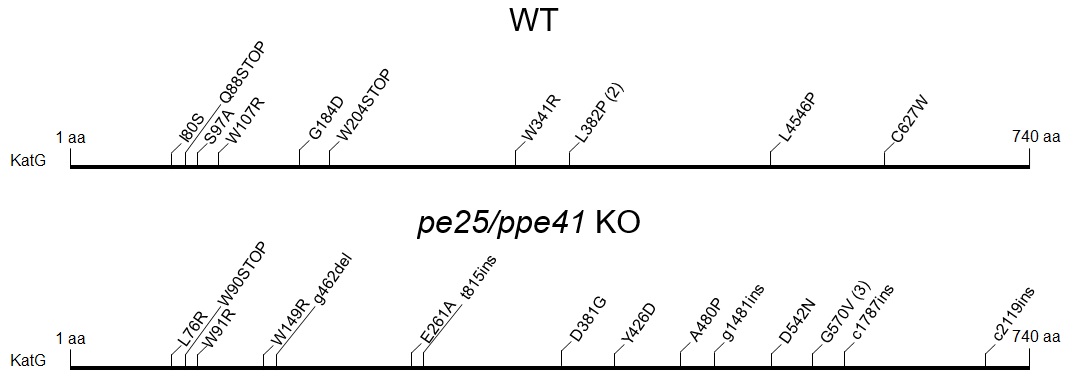
